# Impact of tRNA-induced proline-to-serine mistranslation on the transcriptome of *Drosophila melanogaster*

**DOI:** 10.1101/2024.05.08.593249

**Authors:** Joshua R. Isaacson, Matthew D. Berg, William Yeung, Judit Villén, Christopher J. Brandl, Amanda J. Moehring

## Abstract

Mistranslation is the misincorporation of an amino acid into a polypeptide. Mistranslation has diverse effects on multicellular eukaryotes and is implicated in several human diseases. In *Drosophila melanogaster*, a serine transfer RNA (tRNA) that misincorporates serine at proline codons (P→S) affects male and female flies differently. The mechanisms behind this discrepancy are currently unknown. Here, we compare the transcriptional response of male and female flies to P→S mistranslation to identify genes and cellular processes that underlie sex-specific differences. Both males and females downregulate genes associated with various metabolic processes in response to P→S mistranslation. Males downregulate genes associated with extracellular matrix organization and response to negative stimuli such as wounding, whereas females downregulate aerobic respiration and ATP synthesis genes. Both sexes upregulate genes associated with gametogenesis, but females also upregulate cell cycle and DNA repair genes. These observed differences in the transcriptional response of male and female flies to P→S mistranslation have important implications for the sex-specific impact of mistranslation on disease and tRNA therapeutics.

**ARTICLE SUMMARY:** Proline-to-serine mistranslation affects male and female flies differently, but the mechanisms underlying this discrepancy are unknown. We present a transcriptomic analysis of male and female flies showing that mistranslation disrupts metabolic pathways and gametogenesis in both sexes, whereas processes such as DNA repair and cell cycle regulation are affected only in one sex. This is the first analysis that characterizes sex-specific effects of mistranslation and provides intriguing avenues for future research to understand how mistranslation affects males and females.

## INTRODUCTION

Accurate and efficient translation of mRNA into proteins is required for correct cell function and organism development. Errors during translation can decrease lifespan, induce neurodegeneration, and cause behavioural issues and developmental defects (Lee *et al*. 2006; Lu *et al*. 2014; Reverendo *et al*. 2014; Liu *et al*. 2014). Transfer RNAs (tRNAs) play a major role in determining the fidelity of translation, as do aminoacyl-tRNA-synthetases (aaRSs) which aminoacylate tRNAs with their corresponding amino acid (reviewed in Pang *et al*. 2014). Aminoacyl-tRNA-synthetases recognize specific bases, base pairs, or motifs in their cognate tRNAs to ensure accurate aminoacylation (Hou and Schimmel 1988; Francklyn and Schimmel 1989; Normanly *et al*. 1992; Xue *et al*. 1993; Larkin *et al*. 2002). Since tRNA decoding potential is determined by the anticodon (the nucleotides at positions 34–36 of the tRNA that base-pair with mRNA codons), the anticodon is an identity element for many tRNAs (Schulman and Pelka 1989; Ruff *et al*. 1991; Jahn *et al*. 1991; Tamura *et al*. 1992; Kholod *et al*. 1997; Giegé *et al*. 1998; Zamudio and José 2018; Giegé and Eriani 2023). However, some aaRSs do not use the anticodon to recognize their cognate tRNA. For example, tRNA^Ser^ and tRNA^Ala^ are recognized through an elongated variable arm and a G3:U70 base pair, respectively (McClain and Foss 1988; Francklyn and Schimmel 1989; Achsel and Gross 1993). Because of this, anticodon mutations in tRNA^Ser^ or tRNA^Ala^ genes cause the tRNA to decode noncognate mRNA codons and misincorporate serine or alanine in place of the amino acid normally specified by that codon. This error leads to mistranslation: the incorporation of an amino acid not specified by the standard genetic code. Mistranslation normally occurs at a rate of once per 10^3^–10^6^ codons (Joshi *et al*. 2019; Mordret *et al*. 2019), but tRNA variants or mutant aaRSs can dramatically increase mistranslation (Zimmerman *et al*. 2018; Berg *et al*. 2019b; Zhang *et al*. 2021).

Humans have ∼66 tRNA variants per person, some of which cause mistranslation (Berg *et al*. 2019a; Lant *et al*. 2021; Hasan *et al*. 2023; Davey-Young *et al*. 2024). Mistranslation induces aberrant phenotypes in a variety of organisms, including slow growth in yeast, deformities and decreased lifespan in flies, and cardiac abnormalities and neurodegeneration in mice (Lu *et al*. 2014; Liu *et al*. 2014; Berg *et al*. 2019b, 2021a; Isaacson *et al*. 2022). Previous work in *Saccharomyces cerevisiae* demonstrated that mistranslation affects various biological processes, including translation, stress response, carbohydrate metabolism, and DNA replication (Paredes *et al*. 2012; Berg *et al*. 2021b). The impact of mistranslation is likely more complex in multicellular organisms as codon usage and gene expression vary by tissue and developmental stage (Moriyama and Powell 1997; Dittmar *et al*. 2006; Vicario *et al*. 2008; Allen *et al*. 2022). Transient expression of mistranslating serine tRNA variants in zebrafish embryos upregulated stress response and DNA repair pathways (Reverendo *et al*. 2014), whereas transfection of human cells with mistranslating tRNAs upregulated protein-folding and small molecule catabolism genes (Hou *et al*. 2024). Some mistranslating tRNA variants reduce overall protein synthesis (Lant *et al*. 2021) and alter expression of other tRNAs (Hou *et al*. 2024). Not surprisingly, mistranslating tRNAs have been linked to disease (Goto *et al*. 1990; Shoffner *et al*. 1990; reviewed in Abbott *et al*. 2014; Lant *et al*. 2019).

Sex is an understudied but important influence on organismal response to mistranslation, as male and female physiology differ dramatically due to different metabolic and reproductive requirements (reviewed in Millington and Rideout 2018). Supporting this idea, we previously found that a tRNA^Ser^ variant, which causes proline-to-serine (P→S) mistranslation, increased morphological defects and impaired climbing performance in female fruit flies more than males (Isaacson *et al*. 2022). The mechanisms underlying this difference in male and female response to mistranslation are unknown. The goal of this work is to characterize the impact of P→S mistranslation on the transcriptome of male and female *Drosophila melanogaster* to identify and compare genes and cellular processes that are disrupted in one or both sexes. Using a fly line containing a serine tRNA variant (tRNA^Ser^ ) that induces P→S mistranslation, we found male mistranslating flies primarily downregulate metabolic, developmental, and extracellular matrix organization genes, and upregulate genes associated with spermatogenesis. Female mistranslating flies downregulate genes associated with metabolism and ATP synthesis, and upregulate genes associated with gametogenesis, cell cycle regulation, and DNA repair. As tRNA variants influence disease and are also being assessed as possible therapeutics (reviewed in Anastassiadis and Köhrer 2023 and Coller and Ignatova 2024; Hou *et al*. 2024), it is vital to understand differences in how males and females respond to mistranslating tRNA variants.

## METHODS

### Fly stocks and husbandry

All fly stocks were maintained on standard Bloomington recipe food medium (Bloomington *Drosophila* Stock Center; Bloomington, Indiana) under a 14:10 light:dark cycle at 24° and 70% relative humidity. The tRNA insertion lines used in this study were the same as those described in Isaacson *et al*. (2022). Two fly lines were used: a line containing the wild-type tRNA^Ser^_UGA_ and a line containing the P→S mistranslating tRNA^Ser^ (Isaacson *et al*. 2022). The lines have the same genetic background, and only differ in the type of tRNA transgene that was inserted. The genotype of both lines is as follows:

*w^1118^*; *P{CaryP}attP40[v^+^=tRNA]*/*CyO*, *P{w^+^mC=2xTb^1^-RFP}CyO*; *MKRS*/*TM6B*, *Tb^1^*. Note that the lines used in this study are heterozygous for the inserted tRNA. The *attP40* landing site was selected as it is relatively inert while allowing for strong expression of transgenes (Markstein *et al*. 2008).

### RNA extraction, library preparation, and sequencing

Adult, virgin flies were aged 1–3 days and separated by sex. Ten flies were aspirated into a vial and flash-frozen using liquid nitrogen. Males and females from the tRNA^Ser^ and tRNA^Ser^_UGG, G26A_ (P→S) lines were collected and processed at the same time. Three replicates were collected in this manner for each genotype. RNA was extracted from fly tissue using the protocol outlined in Allen (2016), though volumes of all reagents were halved to account for using less tissue than the protocol specified. Following TRIzol extraction, RNA was measured in a Nanophotometer P300 (Implen, Inc.) and concentration, 260/280 ratio, and 260/230 ratio recorded to assess purity (Supplementary Table S1). To ensure RNA was free of genomic DNA, the remaining 25 µL of RNA was treated with dsDNAse (New England Biolabs Inc) for 30 minutes at 37°. RNA was recovered through a second TRIzol extraction and samples were assessed again using the Nanophotometer to ensure the RNA remained pure. Up to 20 µg of RNA was loaded into RNA-stabilizing tubes, vacuum-dried, and shipped to GeneWiz (South Plainfield, NJ, USA) for total RNA sequencing. If the total amount of RNA was less than 20 µg, then the entire sample was sequenced. Illumina HiSeq 2×150 bp RNA libraries with polyA selection were prepared from each sample. Number of raw reads obtained from each sample ranged from 12.7 million to 68.7 million.

### RNA sequence data processing

Analysis of RNA sequencing data was performed using similar methods to those described in Berg *et al*. (2021b). Short and/or low-quality reads were filtered out using a custom bioinformatics pipeline that utilized Trimmomatic v0.39 (Bolger *et al*. 2014) and FASTQC v0.11.9 (Andrews 2010) to produce filtered libraries containing 8.4–35.4 million reads per sample. Reads were aligned to the *Drosophila melanogaster* reference genome (release r6.41_FB2021_04, downloaded from FlyBase.org, Gramates *et al*. 2022) using STAR v2.7.9a (Dobin *et al*. 2013). Read count data for each gene was summarized using featureCounts v2.0.0 (Liao *et al*. 2014). Only protein-coding genes were included in further analysis. List of protein-coding genes was based on the fly genome assembly BDGP6.46 (Celniker *et al*. 2002; Celniker and Rubin 2003). Parameters and commands used for this pipeline can be found in the extended methods section of supplemental file S2.

### Gene expression and gene ontology analysis

Statistical tests, principal component analysis (PCA), and RNA-Seq data analyses were conducted using R Studio v1.2.5001. RNA-sequencing sample normalization and differential gene expression analysis was performed using the DESeq2 R package (v1.26.0, Love *et al*. 2014), with a Benjamini-Hochberg false discovery rate (FDR) *P*-value cutoff < 0.05. To control for the batch effect identified by the PCA, we specified sample collection day as a covariate in the statistical model fit by ComBat-seq (Zhang *et al*. 2020). Analysis of differentially-expressed genes was performed using WebGestalt’s 2024 release (Liao *et al*. 2019). Lists of down- or upregulated genes were processed by ViSEAGO to produce Gene Ontology (GO) term heatmaps clustered by semantic similarity using Wang’s method (Wang *et al*. 2007; Brionne *et al*. 2019; Gene Ontology Consortium *et al*. 2023). Significantly enriched GO terms were identified by ViSEAGO using the “weight01” algorithm and assessed with Fisher’s exact test. Background gene lists composed of all genes with non-zero read counts for a given sample set (e.g. all female tRNA^Ser^_UGA_ and tRNA^Ser^_UGG, G26A_ samples) were provided to WebGestalt and ViSEAGO during enrichment analysis of that sample set as recommended by Timmons *et al*. (2015) and Wijesooriya *et al*. (2022). Figures were produced using RStudio and Inkscape v1.0.1.

### Validation of RNA-Sequencing results using RT-qPCR

RNA from three new replicates of ten male or female virgin flies containing tRNA^Ser^_UGA_ or tRNA^Ser^ (P→S) aged 1–3 days were extracted using the protocol described above. cDNA was synthesized from RNA using a Maxima H-First Strand cDNA Synthesis Kit (Thermo Scientific™). Quantitative PCRs were performed on three independent replicates in duplicate using 10 ng/µL cDNA template, 500 ng/µL primers, and 1x PowerUp™ SYBR™ Green Master Mix for qPCR (Applied Biosystems™) in a Bio-Rad CFX96 Touch Real-Time PCR Detection System (Bio-Rad Laboratories, Inc.). The Ct values of experimental genes were compared to the Ct values of αTub84B (FBgn0003884) for normalization and statistical analysis, which was performed by the Bio-Rad CFX Manager 3.0 software (Bio-Rad Laboratories, Inc.). A full list of qPCR primers can be found in Supplemental Table S2.

### Clustering analysis

Clustering analysis was performed on the relative fold change of gene expression for tRNA^Ser^_UGG, G26A_ (P→S) compared to control tRNA^Ser^ lines and the relative fold change of gene expression between treatment lines and controls within the microarray data described in Zhou *et al*. (2012). Normalized count data were obtained for all samples and relative fold change compared to controls were calculated for each gene within each treatment. Duplicate genes, genes with < 10 normalized reads, or genes with a relative fold change > |5| were excluded from analysis as Z-score transformation is sensitive to outliers. Relative expression fold change values within each sample were Z-transformed and clustered using the ComplexHeatmap package in RStudio (Gu *et al*. 2016) using Ward’s method (Ward 1963). Male and female data were clustered separately.

## RESULTS

### Identifying mistranslation-induced differentially expressed genes

To analyze the transcriptomic response to serine mistranslation at proline codons in *Drosophila melanogaster*, we sequenced polyA-enriched RNA from 1–3 day old virgin adult male and female flies containing a single copy of either a wild-type tRNA^Ser^ or a tRNA^Ser^ _G26A_ variant that mistranslates proline to serine at a frequency of ∼0.6% per codon (Isaacson *et al*. 2022). The secondary G26A mutation was included in the mistranslating tRNA^Ser^ variant as it disrupts a key modification in tRNA^Ser^ species, reducing mistranslation to survivable levels based on work in yeast (Berg *et al*. 2021a; Boccaletto *et al*. 2022) and flies (Isaaacson *et al*. 2022). tRNA insertion lines were maintained as heterozygotes because naturally-occurring mistranslating tRNA variants are likely to arise as single alleles. Principal component analysis (PCA) was performed on the male and female tRNA^Ser^ and tRNA^Ser^ (P→S) transcriptomic data (Supplemental Figure S1). The first two principal components (PC1 and PC2) summarize ∼55% of the variance of both male and female data. The variation in PC1 captures the batch effect related to the day each sample was collected, as RNA from replicate 1 was harvested a day before replicates 2 and 3. Samples belonging to tRNA^Ser^ or tRNA^Ser^ _G26A_ (P→S) cluster together along the PC2 axis, indicating that the variance explained by PC2 likely represents differences due to the mistranslating tRNA (Figure S1A, B). We corrected the batch effect using ComBat-seq (Zhang *et al*. 2020), and the resulting PCA plots show that samples cluster well and PC1, which represents presence of mistranslation, explains 30–35% of the variance in the data (Figure S1 C, D).

Differentially expressed genes between tRNA^Ser^_UGA_ and tRNA^Ser^ (P→S) were identified using the R package DESeq2 (Love *et al*. 2014). Male and female samples were analysed separately to determine the effects of tRNA^Ser^ (P→S) on each sex. We evaluated 13202 genes with non-zero total read counts in male samples, whereas 12893 genes were evaluated in female samples. MA plots constructed from male or female RNA-sequencing data show that the majority of genes have a log_2_ fold change near zero, as expected (Figure 1A, B). RNA sequencing revealed substantial sex-specific alterations to gene expression in response to mistranslation, as 426 genes were downregulated and 566 genes were upregulated uniquely in males, whereas 507 genes were downregulated and 432 genes upregulated uniquely in females (Wald test performed by DEseq2, Benjamini-Hochberg adjusted *P*-values < 0.05, Figure 1C, D). Only 20 genes were upregulated in both male and female flies containing tRNA^Ser^ (P→S) (Figure 1C), whereas 340 genes were downregulated in both sexes in the mistranslating line (Figure 1D). As shown in Figure 1E, the relative expression of many of the differentially expressed genes differed substantially between the sexes. These results show that P→S mistranslation disrupts expression of largely different sets of genes in males and females.

**Figure 1.**
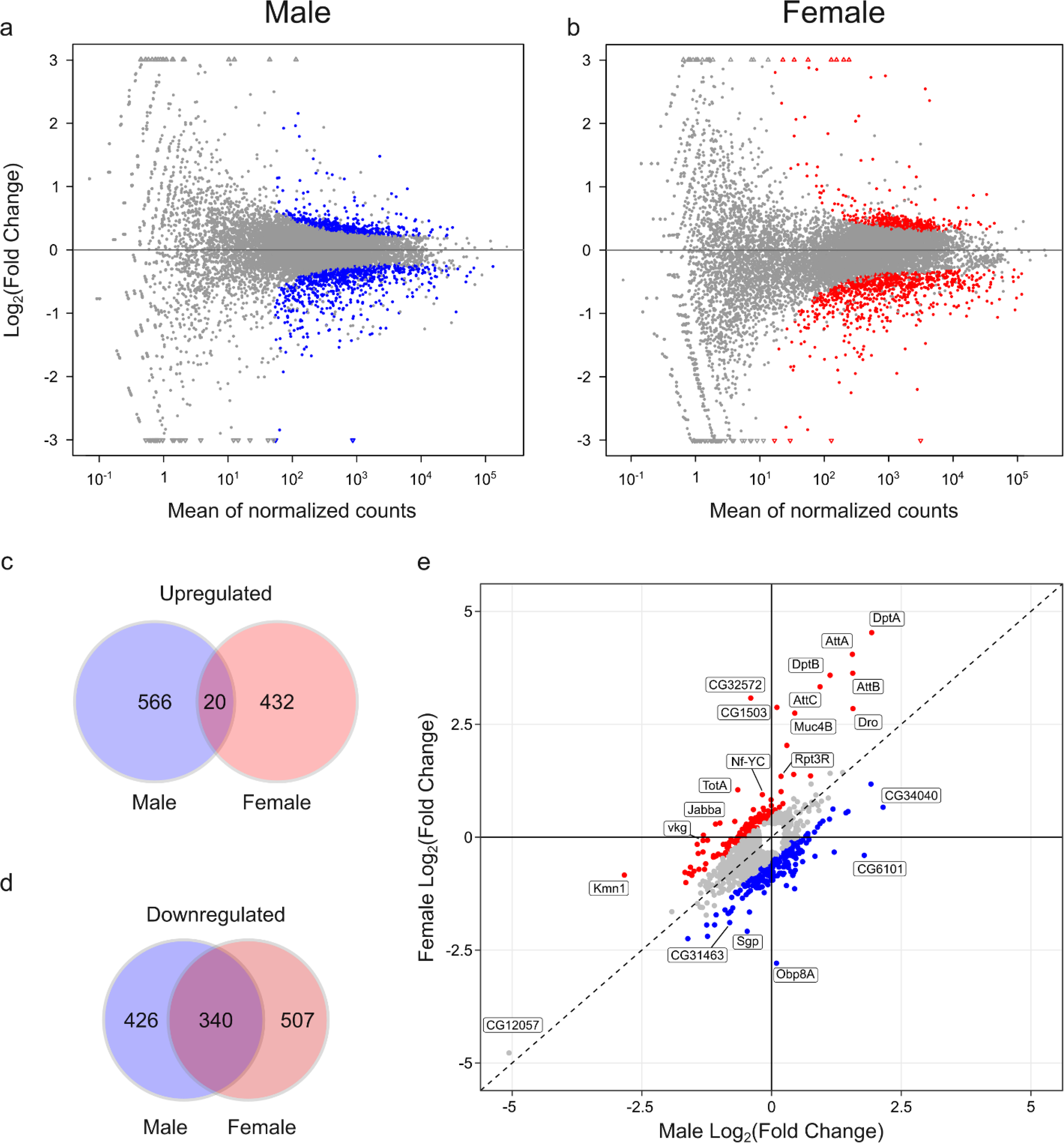
Differentially expressed genes in male or female flies containing tRNA^Ser^ (P→S). **A)** MA plot visualizing the relationship between transcript abundance and the difference in fold change of expression between male tRNA^Ser^ (P→S) and control tRNA^Ser^ samples. Blue points represent genes that are significantly differentially expressed between mistranslating tRNA^Ser^ (P→S) and control tRNA^Ser^ samples, whereas grey points represent genes where the expression change was not statistically significant. Triangular points at the edge of the y-axis indicate genes that have a fold change exceeding the limits of the y-axis. **B)** MA plot visualizing the relationship between transcript abundance and fold change of expression difference between female tRNA^Ser^ (P→S) and control tRNA^Ser^ samples. Red points represent genes that are significantly differentially expressed between mistranslating tRNA^Ser^_UGG, G26A_ (P→S) and control tRNA^Ser^ samples. **C)** Venn diagram showing the number of significantly upregulated (FDR adjusted *P* < 0.05) genes unique to tRNA^Ser^ (P→S) males, females, or genes upregulated in both sexes. **D)** Venn diagram showing the number of significantly downregulated (FDR adjusted *P* < 0.05) genes unique to tRNA^Ser^ (P→S) males, females, or genes downregulated in both sexes. **E)** Scatterplot showing male *vs.* female relative expression for all 1705 genes that were identified as differentially expressed and not filtered out from analysis in either sex. Blue points represent genes that have higher relative expression in mistranslating males compared to females (log_2_ fold change difference > 0.5); red points represent genes with higher relative expression in mistranslating females compared to males. Genes that demonstrate highly sexually-dimorphic patterns of relative expression (log_2_ fold change difference > 1) in response to tRNA^Ser^ (P→S) are labelled. *CG12057* is also labelled due to its strong downregulation in both sexes. The dashed line represents identical fold changes in expression for both males and females.

To provide further support for the transcriptomic data, we confirmed differential expression of six genes using RT-qPCR with RNA extracted from three independent replicates of both male and female flies. We analyzed three genes that were downregulated in both sexes (*CG12057, CG11911, and fiz*), one gene that was upregulated in both sexes (*CG4650*), one gene significantly upregulated in males (*Pif1A*), and one gene that was differentially expressed between males and females (*CG1503*). All genes showed the same pattern of expression in both qPCR and RNA-sequencing analysis for both sexes except for *CG11911*, where the difference between flies containing tRNA^Ser^ or tRNA^Ser^ (P→S) was nonsignificant (Supplemental Figure S2). This rate of non-concordance matches the non-concordance rate of 15–19% between RNA-sequencing and RT-qPCR analysis observed by Everaert *et al*. (2017), who also found non-concordance was more common for short one-exon genes such as *CG11911*.

### Proline-to-serine mistranslation causes sex-specific transcriptional responses

We analysed the lists of differentially-expressed genes using two different tools to identify cellular processes affected by the presence of tRNA^Ser^ (P→S). WebGestalt (Liao *et al*. 2019) was used to identify the ten most enriched Gene Ontology (GO) terms in the list of genes affected by tRNA^Ser^ (P→S). We also used ViSEAGO (Brionne *et al*. 2019) to construct a heatmap of enriched (GO) terms for males and females, allowing for visualization of sex differences in the fly response to P→S mistranslation. All enriched GO terms, their associated *P*-values, and the genes identified in our analysis that belong to those categories are reported in Supplemental file S1. The list of GO terms produced by WebGestalt showed similarities and differences between male and female response to P→S mistranslation. Both males and females downregulated various metabolic processes (Figure 2A), with females primarily downregulating aerobic respiration (e.g., *ox*, *ND-23*, *ND-24*, *UQCR-6.4*, *Cyt-C1*, *COX4*) and males downregulating lipid and fatty acid metabolism (e.g., *Lip4*, *Lsd-1*, *Hacl, FASN1*, *CDase*).

**Figure 2.**
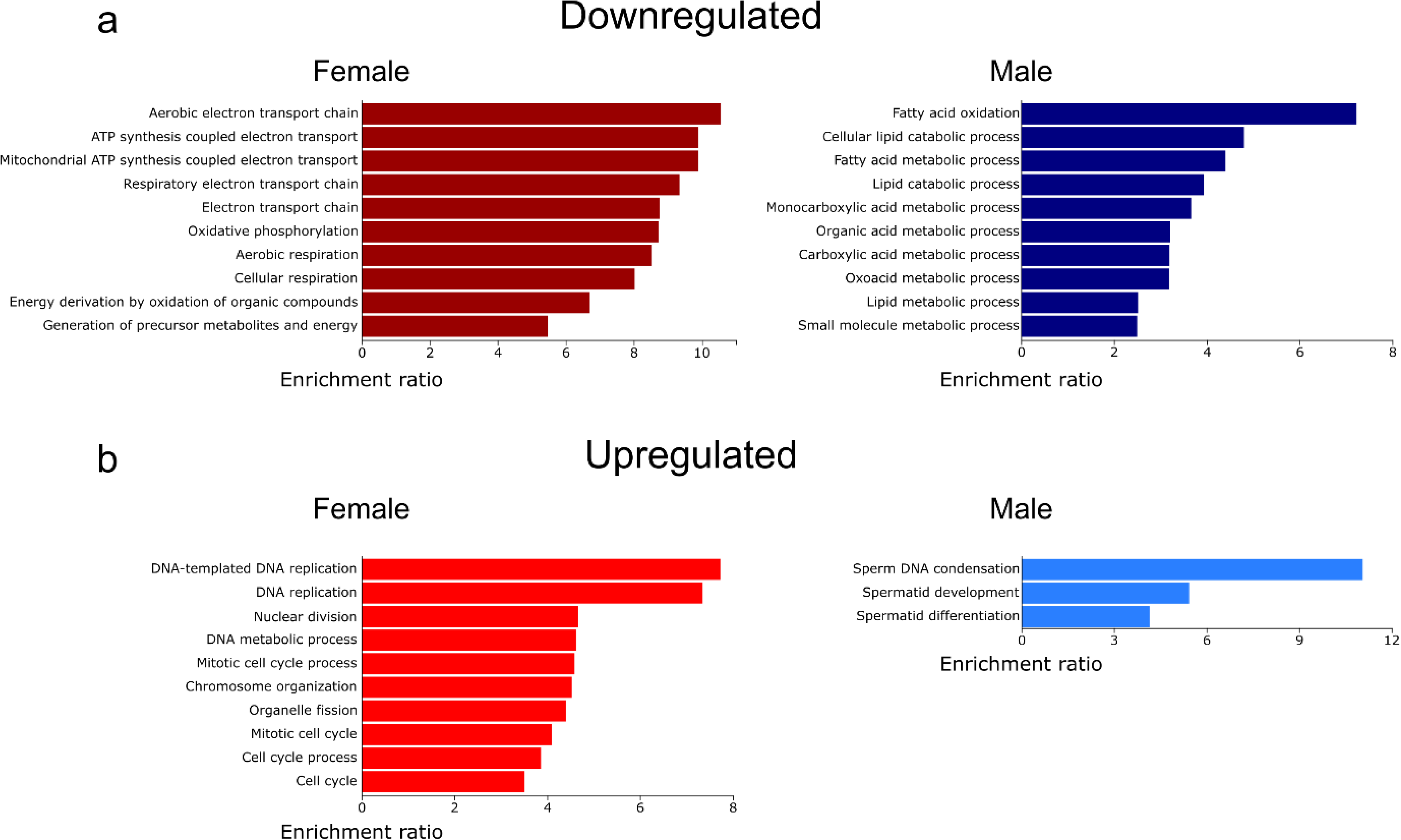
The top 10 significantly enriched gene ontology (GO) terms in the list of genes **A)** downregulated or **B)** upregulated in male or female flies containing tRNA^Ser^_UGG, G26A_ (P→S) compared to control tRNA^Ser^_UGA_ flies. Higher enrichment ratios indicate that the set of genes associated with that GO term were more highly represented in our gene set. Note the differences in scale. Lists were produced using WebGestalt (Liao *et al*. 2019). A list of significantly enriched GO terms and associated statistics is found in Supplemental file S1.

There was limited overlap in the biological processes enriched in the upregulated genes shared between males and females, consistent with our observation that relatively few genes were upregulated in both sexes (Figure 2B). Females upregulated genes associated with cell cycle regulation and cell division (e.g. *CycA*, *CycB*, *Cdc16, APC7*, and *Mink*) as well as genes involved in response to DNA replication (e.g., *DNAlig1*, *PolA1*, *Prim1, RecQ4*, and *Fen1*). Only three biological processes were significantly enriched in the list of upregulated genes in male tRNA^Ser^_UGG, G26A_ (P→S) flies. This may arise because most upregulated genes (438 of 586) are uncharacterized (Supplemental file S1). The three enriched male terms all correspond to male gamete generation and development (e.g., *fan*, *ProtA*, *Pif1A*, and *ntc*).

We next used ViSEAGO to construct heatmaps of GO terms enriched in the set of genes down- or upregulated gene in tRNA^Ser^ (P→S). ViSEAGO clusters GO terms by semantic similarity, so GO terms corresponding to similar biological processes are near each other in the dendrogram (Brionne *et al*. 2019). Functional enrichment was determined using the Fisher’s exact test. Figure 3A further emphasizes the downregulation of genes involved in metabolic processes in response to P→S mistranslation, with different aspects of metabolism being affected in each sex (Figure 3A). In agreement with the WebGestalt results, females downregulated genes associated with oxidative phosphorylation and ATP synthesis whereas males downregulated genes involved in fatty acid and carboxylic acid catabolism. In addition, both males and females downregulated genes involved in chemical or ion transport (e.g., *nrv2*, *blw*, *rumpel*, *snu*). In contrast, biological processes such as response to negative stimuli like wounding (e.g., *Atg2*, *PPO2*, *Hml*, *Tg*) and extracellular structure organization (*Cad99C, LanA, LanB1*, *LanB2, Col4a1, vkg*) were downregulated only in males. Females uniquely downregulated genes associated with muscle function and development, such as myosin (*Mhc, Mlc1, Mlc2*), troponin (*up*, *wupA*), and tropomyosin (*Tm1* and *Tm2*) genes.

**Figure 3.**
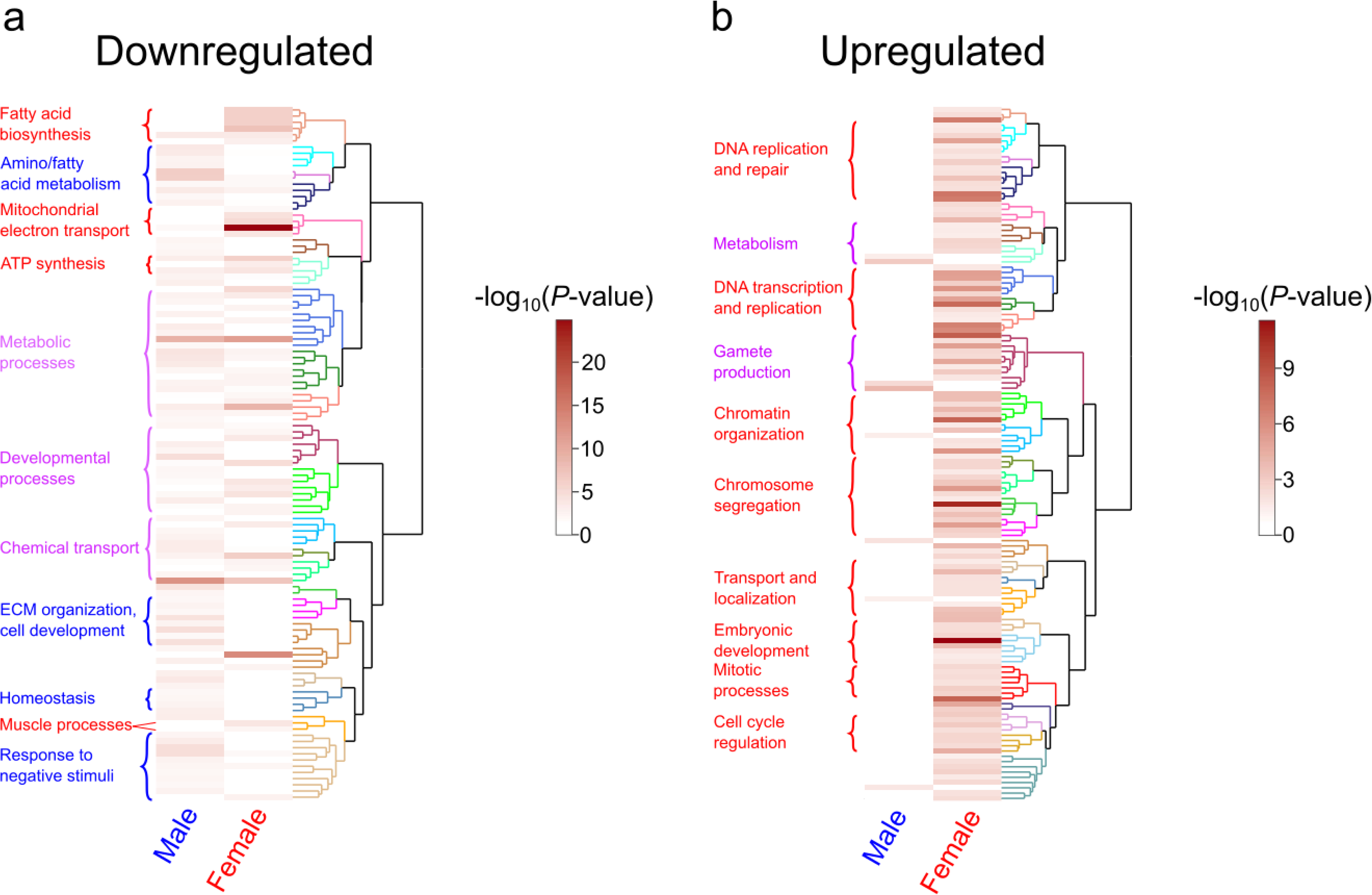
Heatmap of enriched gene ontology (GO) terms from the differentially-expressed genes in male or female flies containing tRNA^Ser^ (P→S). **A)** Heatmap of enriched GO terms in the list of downregulated genes in male and female flies containing tRNA^Ser^ (P→S). Each horizontal bar represents a GO term identified as significantly enriched in male and/or female data. GO terms were clustered by semantic similarity according to ViSEAGO using Wang’s method (Wang *et al*. 2007; Brionne *et al*. 2019). Dendrogram clades of the same color represent semantically similar GO terms. Darker bars within the heatmap represent lower *P*-values as determined through Fisher’s exact test. Notable groups of enriched processes are labelled in blue if enriched in males, red if enriched in females, or purple if enriched in both sexes. **B)** Same as **A)** but using the list of upregulated genes. A full list of enriched GO terms and their associated genes can be found in Supplemental File S1.

When examining the lists of genes upregulated in male or female flies containing tRNA^Ser^_UGG, G26A_ (P→S), ViSEAGO did not identify any GO terms that were significantly enriched in both males and females though we note that gametogenesis and metabolic processes were affected in both sexes (Figure 3B). Of the upregulated genes with identified function, only genes associated with spermatogenesis, protein localization to microtubules, the electron transport chain, and maltose metabolism were enriched in males. For females, in addition to genes associated with cell cycle regulation and DNA repair (discussed above), genes associated with protein and mRNA localization (e.g., *Nup154*, *Elys*, *Fmr1*), development (e.g., *glu*, *mor,* and *fz*), and regulation of gene expression (e.g., *bcd*, *Marf1*, *pum*) were upregulated. Genes involved in antibacterial immune response (e.g., *DptA*, *Dro*, *AttA*, and *BomS5*) were also upregulated in females but not males. These results emphasize that the cellular response to P→S mistranslation differs between male and female flies, and that the difference is particularly pronounced when comparing upregulated genes.

### tRNA-induced P→S mistranslation clusters with heat shock and nutrient stress

Clustering analysis groups genes or treatments based on similarity and is useful to predict functions of uncharacterized genes or identify treatments that produce similar cellular effects (reviewed in Oyelade *et al*. 2016). To identify which environmental or physiological conditions resemble tRNA-induced P→S mistranslation in flies, we clustered the gene expression data from male and female flies containing tRNA^Ser^ with the microarray gene expression data from (Zhou *et al*. 2012), containing the transcriptional response of male and female flies from the same genetic background exposed to 20 different nutritional, chemical, and physiological conditions (Figure 4).

**Figure 4.**
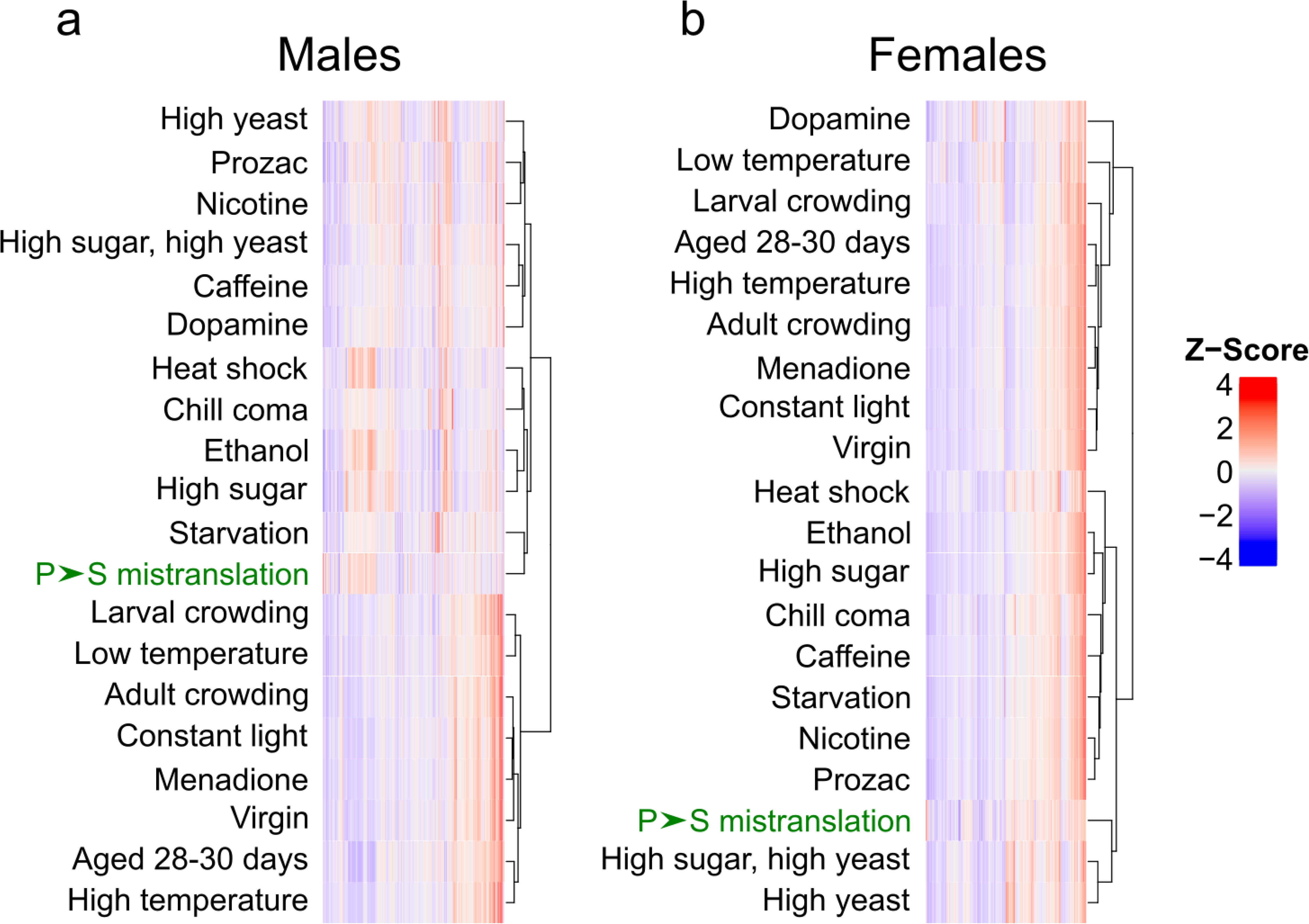
Clustering proline-to-serine mistranslation-induced transcriptome changes with transcriptome changes due to various other physiological or environmental conditions. **A)** Z-score normalized gene expression changes in tRNA^Ser^_UGG, G26A_ (P→S) males relative to tRNA^Ser^_UGA_ (wild-type) males clustered with normalized male gene expression changes from Zhou *et al*. (2012). Genes with fewer than 10 normalized reads or fold changes > |5| for any condition were excluded from analysis. Clustering was performed using the “ComplexHeatmap” R package using Ward’s method (Ward 1963; Gu *et al*. 2016). The P→S mistranslation condition is highlighted in green. **B)** same as **A)** but clustering female data.

As the transcriptomic data acquisition method differed between this study and Zhou *et al*. (2012), we used Z-transformed relative fold changes to compare these datasets. Clustering analysis of male data revealed that tRNA-induced P→S mistranslation induced a transcriptional response most resembling starvation (Figure 4A). Mistranslating males also clustered with temperature or chemical stressors such as heat shock, chill coma, and ethanol exposure. In females, the transcriptional response of tRNA-induced P→S mistranslation most resembled flies reared on high yeast or high sugar and high yeast diets (Figure 4B). Both male and female flies containing tRNA^Ser^ (P→S) clustered with treatments affecting nutrition, which aligns with our observations that various metabolic processes are affected by tRNA^Ser^ (Figure 2A).

## DISCUSSION

### Proline-to-serine mistranslation exerts sex-specific transcriptomic effects

In this study, we examined how *Drosophila melanogaster* males and females alter their transcriptome when exposed to a mistranslating tRNA^Ser^ variant that causes P→S mistranslation. While some biological processes such as carboxylic acid metabolism, chemical transport, and germ cell production were affected in both sexes, we observed a disparity between male and female transcriptional response to P→S mistranslation. This result is consistent with the different physiological and nutritional requirements of male and female flies. Female flies are larger, require a greater quantity and variety of nutrients, and store more triglycerides and glycogen than male flies (Bakker 1959; Wu *et al*. 2020, reviewed in Millington and Rideout 2018;). These requirements are largely due to the increased cost of gamete production in females, which also affects virgin flies as they still devote resources to egg production and laying (Partridge *et al*. 1986; Wu *et al*. 2020). Disruptions to proteostasis, such as mistranslation, would exacerbate this discrepancy between males and females, as maintaining proteostasis requires a substantial proportion of all energy produced by the cell (Buttgereit and Brand 1995; Lahtvee *et al*. 2014). The relatively mild phenotypes previously observed in male flies containing tRNA^Ser^_UGG, G26A_ (P→S) compared to females (Isaacson *et al*. 2022) may in part be due to having more cellular resources available to maintain homeostasis.

One notable group of sex-specific upregulated genes in females were associated with DNA repair and cell cycle regulation. Genes involved with DNA repair are often upregulated in response to cellular stress (Mendez *et al*. 2000; Pregi *et al*. 2017; Sottile and Nadin 2018; Clementi *et al*. 2020). Our observation that DNA repair and cell cycle genes are disrupted in mistranslating flies is consistent with the genetic instability observed by Kalapis *et al*. (2015) in response to mistranslation in yeast. Genetic interactions with mistranslation in yeast and transcriptional responses to mistranslation in human cells also identified the importance of genes involved in cell cycle and DNA damage response (Shcherbakov *et al*. 2019; Berg *et al*. 2021b). Furthermore, mistranslation causes aneuploidy and aberrant nuclear division in yeast species and increases mutation rate in *Escherichia coli* (Balashov and Humayun 2002; Al Mamun *et al*. 2002; Kimata and Yanagida 2004; Silva *et al*. 2007). Mistranslation caused by tRNA^Ser^_UGG, G26A_ may be exerting similar effects in female flies. Interestingly, female flies are less susceptible to sources of DNA damage such as oxidative stress or radiation and are better able to decompose reactive oxygen species than male flies (Parashar *et al*. 2008; Edman *et al*. 2009; Moskalev *et al*. 2011; Niveditha *et al*. 2017). The upregulation of DNA repair genes in mistranslating females may result from their observed increased resistance to stress and DNA damage relative to male flies (reviewed in Pomatto *et al*. 2018). Future studies should examine if flies containing tRNA^Ser^_UGG, G26A_ (P→S) show similar genome instability as mistranslating yeast or *E. coli*.

### Similarity to other transcriptomic studies of tRNA-induced mistranslation

Other studies have examined the transcriptomic effects of tRNA-induced mistranslation on organisms including yeast (Paredes *et al*. 2012; Berg *et al*. 2021b), zebrafish (Reverendo *et al*. 2014), and human cells (HEK293, Hou *et al*. 2024), though none investigated how males and females differ in their response to mistranslation. Paredes *et al*. (2012) engineered a tRNA^Ser^ variant that mistranslates leucine-to-serine in yeast and observed upregulation of stress response chaperone genes and downregulation of protein synthesis. When clustered with various environmental stresses, the mistranslating yeast transcriptome most resembled nutrient stresses such as nitrogen deprivation and amino acid starvation, which agrees with our results in flies. Zebrafish embryos transiently expressing mistranslating tRNA^Ser^ variants similarly downregulate protein synthesis and upregulate stress response genes and genes associated with DNA damage and repair (Reverendo *et al*. 2014). Human cells transfected with mistranslating tRNA^Arg^ variants upregulate genes involved in protein folding and endoplasmic reticulum stress (Hou *et al*. 2024). Interestingly, some mistranslating tRNA^Arg^ variants have minimal effects on the transcriptome. While we did not observe significant downregulation of genes involved in protein synthesis in males or females containing tRNA^Ser^ (P→S), female flies containing tRNA^Ser^ upregulated genes involved in DNA damage and repair, which aligns with the previous studies. Overall, our data is consistent with previous work characterizing the transcriptomic effects of mistranslation in other organisms while uncovering novel sex-specific differences in these general responses.

### Future work and conclusions

These transcriptomic results provide intriguing avenues for future research. *D. melanogaster* tissues have different codon usages and tRNA expression profiles and thus might be differently susceptible to tRNA^Ser^ variants that cause P→S mistranslation (Dittmar *et al*. 2006; Allen *et al*. 2022). A focused transcriptomic approach centered on specific cell types, such as neurons or muscle, could reveal trends that are difficult to observe from whole-fly transcriptomics. Testing other life stages could also reveal stage-specific transcriptomic responses to mistranslating tRNA variants. Different types of mistranslation exert unique cellular effects (Berg *et al*. 2021b; Cozma *et al*. 2023; Hou *et al*. 2024; Davey-Young *et al*. 2024), so testing other amino acid substitutions will uncover which cellular responses are common to mistranslation and which are unique to specific substitutions.

The differentially-expressed genes identified in this analysis can be targeted using available *D. melanogaster* knockout lines to determine which are necessary for the fly response to mistranslation. The uncharacterized gene *CG12057* is worthy of further investigation as its expression was reduced >25-fold in both male and female tRNA^Ser^ (P→S) flies. *CG12057* is primarily expressed in the midgut and its expression is impacted by various stresses, including hypoxia, infection, and mitochondrial dysfunction (Carpenter *et al*. 2009; Fernández-Ayala *et al*. 2010; Mosqueira *et al*. 2010; Moskalev *et al*. 2015; Krause *et al*. 2022). Determining the function of *CG12057* could provide insight into how flies cope with cellular stress. Further investigation into the cellular processes disrupted by P→S mistranslation may elucidate the genetic and physiological mechanisms behind sex-specific response to mistranslation and the striking phenotypes observed in mistranslating adult flies (Isaacson *et al*. 2022). Overall, this study demonstrates that sex strongly affects response to mistranslation and must be considered when studying mistranslation in sexually-dimorphic organisms.

## DATA AVAILIBILITY

Fly lines are available upon request. The authors affirm that all data necessary for confirming the conclusions of the article are present within the article, figures, and supplemental material. A full list of all differentially expressed genes and all significantly enriched GO terms can be found in Supplemental file S1. Supplemental file S2 contains all supplementary figures, tables, and methods. Supplemental file S3 contains R code used to analyze RNA-sequencing data and perform clustering and ViSEAGO analysis. All raw and processed data can be found at the NCBI GEO database using the accession number GSE256332.

## ACKNOWLEDGEMENTS

We would like to thank Dr. Patrick O’Donoghue, Dr. Robert Cumming, and Ecaterina Cozma for their feedback and guidance on the manuscript, as well as Dr. Gregory Gloor for his advice on data analysis and interpretation. This work was supported by Natural Sciences and Engineering Research Council of Canada (NSERC) grants to CJB [RGPIN-2020-07046] and AJM [RGPIN-2020-06464], NIH grants to JV [R01AG056359, R56AG049494 and R35GM119536], as well as a University of Western Ontario Medical & Health Science Research Board seed grant to AJM. JRI and MDB were supported by an NSERC Postgraduate Scholarship (Doctoral) and NSERC Canada Graduate Scholarship (Doctoral), respectively.

---

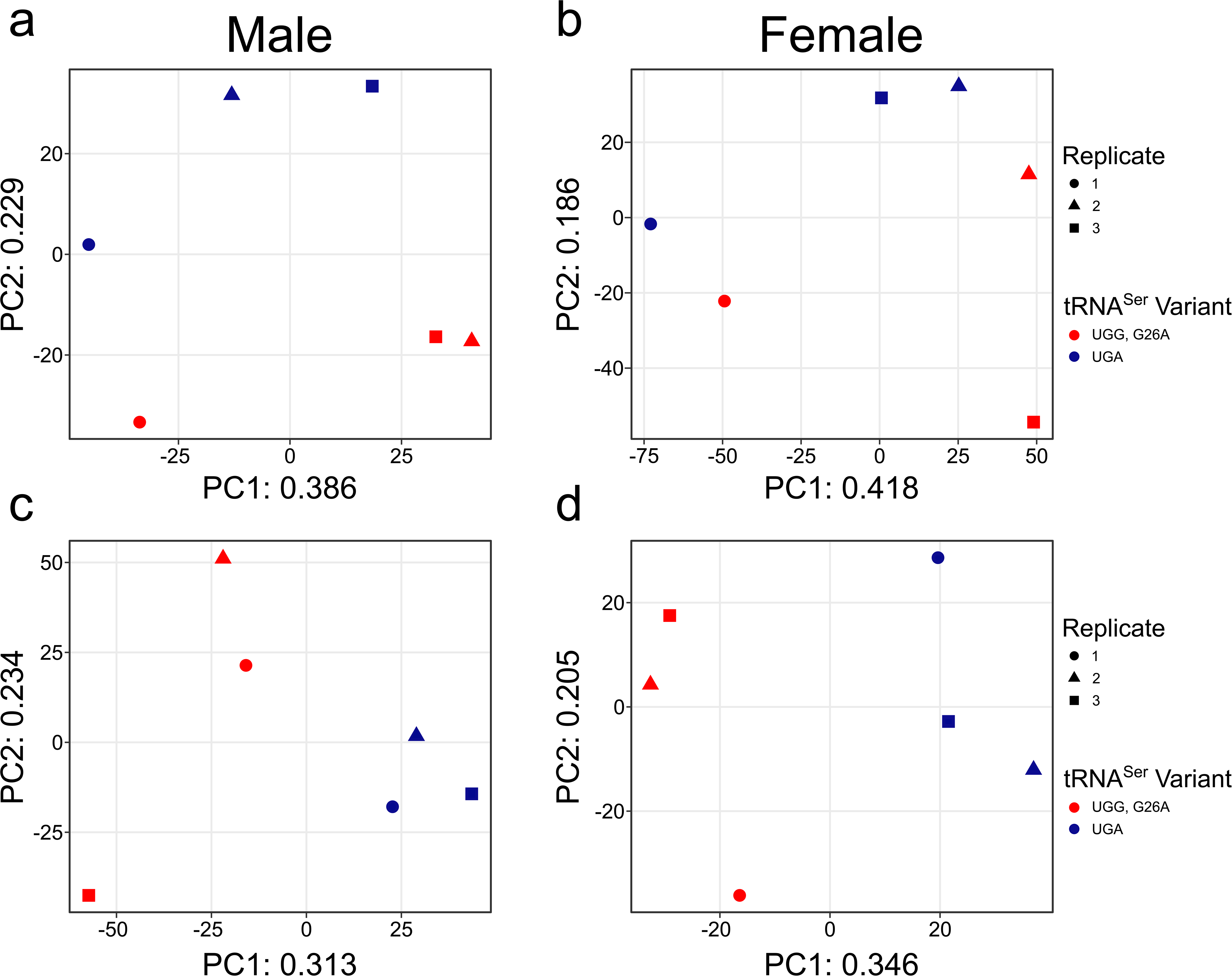

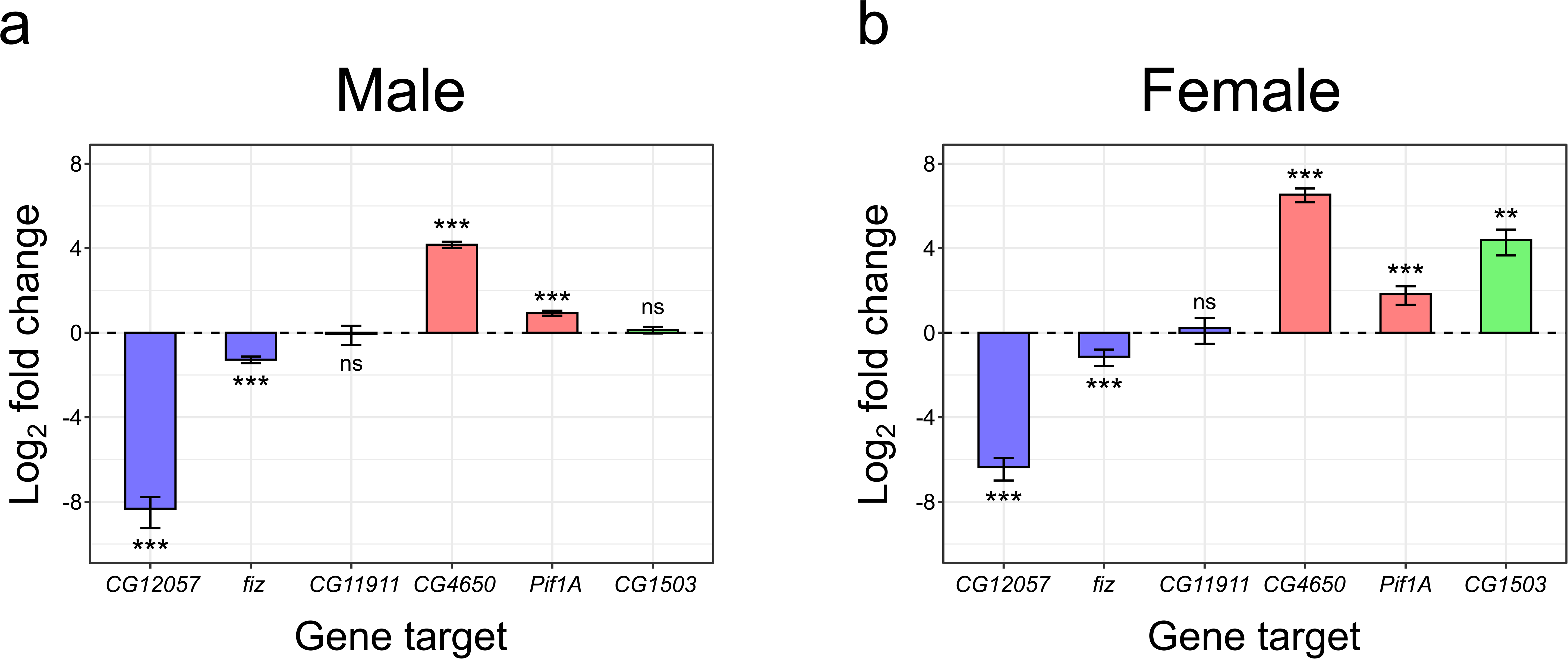

## LITERATURE CITED

Abbott, J. A., C. S. Francklyn, and S. M. Robey-Bond, 2014 Transfer RNA and human disease. Front. Genet. 5: 158.

Achsel, T., and H. J. Gross, 1993 Identity determinants of human tRNA^Ser^: sequence elements necessary for serylation and maturation of a tRNA with a long extra arm. EMBO J. 12: 3333–3338.

Allen, E., 2016 RNA Extraction from *Drosophila* tissues using TRIzol reagent. protocols.io 10.17504/protocols.io.fgtbjwn.

Allen, S. R., R. K. Stewart, M. Rogers, I. J. Ruiz, E. Cohen et al., 2022 Distinct responses to rare codons in select *Drosophila* tissues. eLife 11: e76893.

Anastassiadis, T., and C. Köhrer, 2023 Ushering in the era of tRNA medicines. J. Biol. Chem. 299: 105246.

Andrews, S., 2010 FASTQC: A quality control tool for high throughput sequence data. Bakker, K., 1959 Feeding period, growth, and pupation in larvae of *Drosophila melanogaster*. Entomol. Exp. Appl. 2: 171–186.

Balashov, S., and M. Z. Humayun, 2002 Mistranslation induced by streptomycin provokes a RecABC/RuvABC-dependent mutator phenotype in *Escherichia coli* cells. J. Mol. Biol. 315: 513–527.

Berg, M. D., D. J. Giguere, J. S. Dron, J. T. Lant, J. Genereaux et al., 2019a Targeted sequencing reveals expanded genetic diversity of human transfer RNAs. RNA Biol. 16: 1574–1585.

Berg, M. D., J. R. Isaacson, E. Cozma, J. Genereaux, P. Lajoie et al., 2021a Regulating expression of mistranslating tRNAs by readthrough RNA polymerase II transcription. ACS Synth. Biol. 10: 3177–3189.

Berg, M. D., Y. Zhu, J. Genereaux, B. Y. Ruiz, R. A. Rodriguez-Mias et al., 2019b Modulating mistranslation potential of tRNA^Ser^ in *Saccharomyces cerevisiae*. Genetics 213: 849–863.

Berg, M. D., Y. Zhu, B. Y. Ruiz, R. Loll-Krippleber, J. Isaacson et al., 2021b The amino acid substitution affects cellular response to mistranslation. G3 Genes|Genomes|Genetics 11: jkab218.

Boccaletto, P., F. Stefaniak, A. Ray, A. Cappannini, S. Mukherjee et al., 2022 MODOMICS: a database of RNA modification pathways. 2021 update. Nucleic Acids Res. 50: D231–D235.

Bolger, A. M., M. Lohse, and B. Usadel, 2014 Trimmomatic: a flexible trimmer for Illumina sequence data. Bioinformatics 30: 2114–2120.

Brionne, A., A. Juanchich, and C. Hennequet-Antier, 2019 ViSEAGO: a Bioconductor package for clustering biological functions using Gene Ontology and semantic similarity. BioData Min. 12: 16.

Buttgereit, F., and M. D. Brand, 1995 A hierarchy of ATP-consuming processes in mammalian cells. Biochem. J. 312: 163–167.

Carpenter, J., S. Hutter, J. F. Baines, J. Roller, S. S. Saminadin-Peter et al., 2009 The transcriptional response of *Drosophila melanogaster* to infection with the sigma virus (*Rhabdoviridae*). PLoS One 4: e6838.

Celniker, S. E., and G. M. Rubin, 2003 The *Drosophila Melanogaster* genome. Annu. Rev. Genomics Hum. Genet. 4: 89–117.

Celniker, S. E., D. A. Wheeler, B. Kronmiller, J. W. Carlson, A. Halpern et al., 2002 Finishing a whole-genome shotgun: release 3 of the *Drosophila melanogaster* euchromatic genome sequence. Genome Biol. 3: research0079.1.

Clementi, E., L. Inglin, E. Beebe, C. Gsell, Z. Garajova et al., 2020 Persistent DNA damage triggers activation of the integrated stress response to promote cell survival under nutrient restriction. BMC Biol. 18: 36.

Coller, J., and Z. Ignatova, 2024 tRNA therapeutics for genetic diseases. Nat. Rev. Drug Discov. 23: 108–125.

Cozma, E., M. Rao, M. Dusick, J. Genereaux, R. A. Rodriguez-Mias et al., 2023 Anticodon sequence determines the impact of mistranslating tRNA^Ala^ variants. RNA Biol. 20: 791– 804.

Davey-Young, J., F. Hasan, R. Tennakoon, P. Rozik, H. Moore et al., 2024 Mistranslating the genetic code with leucine in yeast and mammalian cells. RNA Biol. 21: 1–23.

Dittmar, K. A., J. M. Goodenbour, and T. Pan, 2006 Tissue-specific differences in human transfer RNA expression. PLoS Genet. 2: e221.

Dobin, A., C. A. Davis, F. Schlesinger, J. Drenkow, C. Zaleski et al., 2013 STAR: ultrafast universal RNA-seq aligner. Bioinformatics 29: 15–21.

Edman, U., A. M. Garcia, R. A. Busuttil, D. Sorensen, M. Lundell et al., 2009 Lifespan extension by dietary restriction is not linked to protection against somatic DNA damage in *Drosophila melanogaster*. Aging Cell 8: 331–338.

Everaert, C., M. Luypaert, J. L. V. Maag, Q. X. Cheng, M. E. Dinger et al., 2017 Benchmarking of RNA-sequencing analysis workflows using whole-transcriptome RT-qPCR expression data. Sci. Rep. 7: 1559.

Fernández-Ayala, D. J. M., S. Chen, E. Kemppainen, K. M. C. O’Dell, and H. T. Jacobs, 2010 Gene expression in a *Drosophila* model of mitochondrial disease. PLoS One 5: e8549.

Francklyn, C., and P. Schimmel, 1989 Aminoacylation of RNA minihelices with alanine. Nature 337: 478–481.

Gene Ontology Consortium, S. A. Aleksander, J. Balhoff, S. Carbon, J. M. Cherry et al., 2023 The Gene Ontology knowledgebase in 2023. Genetics 224: iyad031.

Giegé, R., and G. Eriani, 2023 The tRNA identity landscape for aminoacylation and beyond. Nucleic Acids Res. 51: 1528–1570.

Giegé, R., M. Sissler, and C. Florentz, 1998 Universal rules and idiosyncratic features in tRNA identity. Nucleic Acids Res. 26: 5017–5035.

Goto, Y. I., I. Nonaka, and S. Horai, 1990 A mutation in the tRNA^Leu(UUR)^ gene associated with the MELAS subgroup of mitochondrial encephalomyopathies. Nature 348: 651–653.

Gramates, L. S., J. Agapite, H. Attrill, B. R. Calvi, M. A. Crosby et al., 2022 FlyBase: a guided tour of highlighted features. Genetics 220:.

Gu, Z., R. Eils, and M. Schlesner, 2016 Complex heatmaps reveal patterns and correlations in multidimensional genomic data. Bioinformatics 32: 2847–2849.

Hasan, F., J. T. Lant, and P. O’Donoghue, 2023 Perseverance of protein homeostasis despite mistranslation of glycine codons with alanine. Philos. Trans. R. Soc. B Biol. Sci. 378: 20220029.

Hou, Y. M., and P. Schimmel, 1988 A simple structural feature is a major determinant of the identity of a transfer RNA. Nature 333: 140–145.

Hou, Y., W. Zhang, P. T. McGilvray, M. Sobczyk, T. Wang et al., 2024 Engineered mischarged transfer RNAs for correcting pathogenic missense mutations. Mol. Ther. 32: 352–371.

Isaacson, J. R., M. D. Berg, B. Charles, J. Jagiello, J. Villén et al., 2022 A novel mistranslating tRNA model in *Drosophila melanogaster* has diverse, sexually dimorphic effects. G3 Genes|Genomes|Genetics 12: jkac035.

Jahn, M., M. J. Rogers, and D. Söll, 1991 Anticodon and acceptor stem nucleotides in tRNA^Gln^ are major recognition elements for *E. coli* glutaminyl-tRNA synthetase. Nature 352: 258– 260.

Joshi, K., L. Cao, and P. J. Farabaugh, 2019 The problem of genetic code misreading during protein synthesis. Yeast 36: 35–42.

Kalapis, D., A. R. Bezerra, Z. Farkas, P. Horvath, Z. Bódi et al., 2015 Evolution of robustness to protein mistranslation by accelerated protein turnover. PLOS Biol. 13: e1002291.

Kholod, N. S., N. V Pan’kova, S. G. Mayorov, A. I. Krutilina, M. G. Shlyapnikov et al., 1997 Transfer RNA^Phe^ isoacceptors possess non-identical set of identity elements at high and low Mg^2+^ concentration. FEBS Lett. 411: 123–127.

Kimata, Y., and M. Yanagida, 2004 Suppression of a mitotic mutant by tRNA-Ala anticodon mutations that produce a dominant defect in late mitosis. J. Cell Sci. 117: 2283–2293.

Krause, S. A., G. Overend, J. A. T. Dow, and D. P. Leader, 2022 FlyAtlas 2 in 2022: enhancements to the *Drosophila melanogaster* expression atlas. Nucleic Acids Res. 50: D1010–D1015.

Lahtvee, P.-J., A. Seiman, L. Arike, K. Adamberg, and R. Vilu, 2014 Protein turnover forms one of the highest maintenance costs in *Lactococcus lactis*. Microbiology 160: 1501–1512.

Lant, J. T., M. D. Berg, I. U. Heinemann, C. J. Brandl, and P. O’Donoghue, 2019 Pathways to disease from natural variations in human cytoplasmic tRNAs. J. Biol. Chem. 294: 5294– 5308.

Lant, J. T., R. Kiri, M. L. Duennwald, and P. O’Donoghue, 2021 Formation and persistence of polyglutamine aggregates in mistranslating cells. Nucleic Acids Res. 49: 11883–11899.

Larkin, D. C., A. M. Williams, S. A. Martinis, and G. E. Fox, 2002 Identification of essential domains for *Escherichia coli* tRNA^Leu^ aminoacylation and amino acid editing using minimalist RNA molecules. Nucleic Acids Res. 30: 2103–2113.

Lee, J. W., K. Beebe, L. A. Nangle, J. Jang, C. M. Longo-Guess et al., 2006 Editing-defective tRNA synthetase causes protein misfolding and neurodegeneration. Nature 443: 50–55.

Liao, Y., G. K. Smyth, and W. Shi, 2014 featureCounts: an efficient general purpose program for assigning sequence reads to genomic features. Bioinformatics 30: 923–930.

Liao, Y., J. Wang, E. J. Jaehnig, Z. Shi, and B. Zhang, 2019 WebGestalt 2019: gene set analysis toolkit with revamped UIs and APIs. Nucleic Acids Res. 47: W199–W205.

Liu, Y., J. S. Satz, M. N. Vo, L. A. Nangle, P. Schimmel et al., 2014 Deficiencies in tRNA synthetase editing activity cause cardioproteinopathy. Proc. Natl. Acad. Sci. USA. 111: 17570–17575.

Love, M. I., W. Huber, and S. Anders, 2014 Moderated estimation of fold change and dispersion for RNA-seq data with DESeq2. Genome Biol. 15: 1–21.

Lu, J., M. Bergert, A. Walther, and B. Suter, 2014 Double-sieving-defective aminoacyl-tRNA synthetase causes protein mistranslation and affects cellular physiology and development. Nat. Commun. 5: 1–13.

Al Mamun, A. A. M., K. J. Marians, and M. Z. Humayun, 2002 DNA polymerase III from *Escherichia coli* cells expressing *mutA* mistranslator tRNA is error-prone. J. Biol. Chem. 277: 46319–46327.

Markstein, M., C. Pitsouli, C. Villalta, S. E. Celniker, and N. Perrimon, 2008 Exploiting position effects and the gypsy retrovirus insulator to engineer precisely expressed transgenes. Nat. Genet. 40: 476–483.

McClain, W. H., and K. Foss, 1988 Changing the identity of a tRNA by introducing a G-U wobble pair near the 3′ acceptor end. Science. 240: 793–796.

Mendez, F., M. Sandigursky, W. A. Franklin, M. K. Kenny, R. Kureekattil et al., 2000 Heat-shock proteins associated with base excision repair enzymes in HeLa cells. Radiat. Res. 153: 186–95.

Millington, J. W., and E. J. Rideout, 2018 Sex differences in *Drosophila* development and physiology. Curr. Opin. Physiol. 6: 46–56.

Mordret, E., O. Dahan, O. Asraf, R. Rak, A. Yehonadav et al., 2019 Systematic setection of amino acid substitutions in proteomes reveals mechanistic basis of ribosome errors and selection for translation fidelity. Mol. Cell 75: 427–441.e5.

Moriyama, E. N., and J. R. Powell, 1997 Codon usage bias and tRNA abundance in Drosophila. J. Mol. Evol. 45: 514–523.

Moskalev, A. A., E. N. Plyusnina, and M. V. Shaposhnikov, 2011 Radiation hormesis and radioadaptive response in *Drosophila melanogaster* flies with different genetic backgrounds: the role of cellular stress-resistance mechanisms. Biogerontology 12: 253– 263.

Moskalev, A., S. Zhikrivetskaya, G. Krasnov, M. Shaposhnikov, E. Proshkina et al., 2015 A comparison of the transcriptome of *Drosophila melanogaster* in response to entomopathogenic fungus, ionizing radiation, starvation and cold shock. BMC Genomics 16: S8.

Mosqueira, M., G. Willmann, H. Ruohola-Baker, and T. S. Khurana, 2010 Chronic hypoxia Iipairs muscle function in the *Drosophila* model of Duchenne’s muscular dystrophy (DMD). PLoS One 5: e13450.

Niveditha, S., S. Deepashree, S. R. Ramesh, and T. Shivanandappa, 2017 Sex differences in oxidative stress resistance in relation to longevity in *Drosophila melanogaster*. J. Comp. Physiol. B 187: 899–909.

Normanly, J., T. Ollick, and J. Abelson, 1992 Eight base changes are sufficient to convert a leucine-inserting tRNA into a serine-inserting tRNA. Proc. Natl. Acad. Sci. USA. 89: 5680– 5684.

Oyelade, J., I. Isewon, F. Oladipupo, O. Aromolaran, E. Uwoghiren et al., 2016 Clustering algorithms: their application to gene expression data. Bioinform. Biol. Insights 10: BBI.S38316.

Pang, Y. L. J., K. Poruri, and S. A. Martinis, 2014 tRNA synthetase: tRNA aminoacylation and beyond. Wiley Interdiscip. Rev. RNA 5: 461–480.

Parashar, V., S. Frankel, A. G. Lurie, and B. Rogina, 2008 The effects of age on radiation resistance and oxidative stress in adult *Drosophila melanogaster*. Radiat. Res. 169: 707– 711.

Paredes, J. A., L. Carreto, J. Simões, A. R. Bezerra, A. C. Gomes et al., 2012 Low level genome mistranslations deregulate the transcriptome and translatome and generate proteotoxic stress in yeast. BMC Biol. 10: 55.

Partridge, L., K. Fowler, S. Trevitt, and W. Sharp, 1986 An examination of the effects of males on the survival and egg-production rates of female *Drosophila melanogaster*. J. Insect Physiol. 32: 925–929.

Pomatto, L. C. D., J. Tower, and K. J. A. Davies, 2018 Sexual dimorphism and aging differentially regulate adaptive homeostasis. J. Gerontol. Ser. A 73: 141–149.

Pregi, N., L. M. Belluscio, B. G. Berardino, D. S. Castillo, and E. T. Cánepa, 2017 Oxidative stress-induced CREB upregulation promotes DNA damage repair prior to neuronal cell death protection. Mol. Cell. Biochem. 425: 9–24.

Reverendo, M., A. R. Soares, P. M. Pereira, L. Carreto, V. Ferreira et al., 2014 tRNA mutations that affect decoding fidelity deregulate development and the proteostasis network in zebrafish. RNA Biol. 11: 1199–1213.

Ruff, M., S. Krishnaswamy, M. Boeglin, A. Poterszman, A. Mitschler et al., 1991 Class II aminoacyl transfer RNA synthetases: crystal structure of yeast aspartyl-tRNA synthetase complexed with tRNA^Asp^. Science 252: 1682–1689.

Schulman, L. H., and H. Pelka, 1989 The anticodon contains a major element of the identity of arginine transfer RNAs. Science 246: 1595–1597.

Shcherbakov, D., Y. Teo, H. Boukari, A. Cortes-Sanchon, M. Mantovani et al., 2019 Ribosomal mistranslation leads to silencing of the unfolded protein response and increased mitochondrial biogenesis. Commun. Biol. 2: 381.

Shoffner, J. M., M. T. Lott, A. M. S. Lezza, P. Seibel, S. W. Ballinger et al., 1990 Myoclonic epilepsy and ragged-red fiber disease (MERRF) is associated with a mitochondrial DNA tRNA^Lys^ mutation. Cell 61: 931–937.

Silva, R. M., J. A. Paredes, G. R. Moura, B. Manadas, T. Lima-Costa et al., 2007 Critical roles for a genetic code alteration in the evolution of the genus *Candida*. EMBO J. 26: 4555– 4565.

Sottile, M. L., and S. B. Nadin, 2018 Heat shock proteins and DNA repair mechanisms: an updated overview. Cell Stress Chaperones 23: 303–315.

Tamura, K., H. Himeno, H. Asahara, T. Hasegawa, and M. Shimizu, 1992 *In vitro* study of *E. coli* tRNA^Arg^ and tRNA^Lys^ identity elements. Nucleic Acids Res. 20: 2335–2339.

Timmons, J. A., K. J. Szkop, and I. J. Gallagher, 2015 Multiple sources of bias confound functional enrichment analysis of global -omics data. Genome Biol. 16: 1–3.

Vicario, S., C. E. Mason, K. P. White, and J. R. Powell, 2008 Developmental stage and level of codon usage bias in *Drosophila*. Mol. Biol. Evol. 25: 2269–2277.

Wang, J. Z., Z. Du, R. Payattakool, P. S. Yu, and C. F. Chen, 2007 A new method to measure the semantic similarity of GO terms. Bioinformatics 23: 1274–1281.

Ward, J. H., 1963 Hierarchical grouping to optimize an objective function. J. Am. Stat. Assoc. 58: 236–244.

Wijesooriya, K., S. A. Jadaan, K. L. Perera, T. Kaur, and M. Ziemann, 2022 Urgent need for consistent standards in functional enrichment analysis. PLOS Comput. Biol. 18: e1009935.

Wu, Q., G. Yu, X. Cheng, Y. Gao, X. Fan et al., 2020 Sexual dimorphism in the nutritional requirement for adult lifespan in *Drosophila melanogaster*. Aging Cell 19:.

Xue, H., W. Shens, R. Giegeq, J. Tze-, and F. Wongii, 1993 Identity elements of tRNA^Trp^. Identification and evolutionary conservation. J. Biol. Chem. 268: 9316–9322.

Zamudio, G. S., and M. V. José, 2018 Identity elements of tRNA as derived from information analysis. Orig. Life Evol. Biosph. 48: 73–81.

Zhang, Y., G. Parmigiani, and W. E. Johnson, 2020 *ComBat-seq*: batch effect adjustment for RNA-seq count data. NAR Genomics Bioinforma. 2:.

Zhang, H., J. Wu, Z. Lyu, and J. Ling, 2021 Impact of alanyl-tRNA synthetase editing deficiency in yeast. Nucleic Acids Res. 49: 9953–9964.

Zhou, S., T. G. Campbell, E. A. Stone, T. F. C. Mackay, and R. R. H. Anholt, 2012 Phenotypic plasticity of the *Drosophila* transcriptome. PLOS Genet. 8: e1002593.

Zimmerman, S. M., Y. Kon, A. C. Hauke, B. Y. Ruiz, S. Fields et al., 2018 Conditional accumulation of toxic tRNAs to cause amino acid misincorporation. Nucleic Acids Res. 46: 7831–7843.

